# Growth, dissolution and segregation of genetically encoded RNA droplets by ribozyme catalysis

**DOI:** 10.1101/2025.08.29.673008

**Authors:** Franziska Giessler, William Verstraeten, Tobias Abele, Stefan J. Maurer, Luca Monari, Kerstin Göpfrich

## Abstract

Active droplets, membraneless compartments driven by internal chemical reactions, are compelling models for protocells and synthetic life. A central challenge is to program their dynamic behaviors using heritable genetic information, which would grant them the capacity to evolve. Here, we create transiently active RNA droplets by integrating sites for ribozyme catalysis directly into the sequence of self-assembling, four-arm RNA nanostars. To enable perfusion and observe the resulting dynamics over time, we develop a method for trapping individual droplets in hydrogel cages by targeted *in situ* photopolymerization. This enables us to quantify the sequence-programmable droplet dissolution and to control the degradation kinetics by choosing between fast (hammerhead) and slow (hairpin) ribozymes. Furthermore, we trigger the segregation of mixed droplet populations via the sequence-specific cleavage of a chimeric linker RNA. The droplet-encapsulated DNA templates code for the regrowth of new droplets, establishing the proof-of-principle for a minimal, genetically encoded cycle of dissolution and regrowth. By directly linking RNA sequence to droplet stability, composition, and life-cycle dynamics, our work provides a robust platform for engineering evolvable materials and advancing the bottom-up construction of synthetic cells.

## Introduction

Active droplets are membraneless compartments with life-like behaviors such as growth, decay, and selforganization driven by internal chemical reactions. They could offer a route to compartmentalize early biochemistry at the origins of life [1] and are compelling models for the forward-looking goal of creating synthetic life within the fields of systems chemistry and bottom-up synthetic biology [2]. Droplets, also termed biomolecular condensates or coacervates, form spontaneously when a solution of macromolecules undergoes liquid-liquid phase separation, creating a dense phase distinct from its dilute environment driven by weak, multivalent interactions [3]. Their ‘active’ nature arises from internal chemical reactions that consume energy to maintain the system out-of-equilibrium [4]. Active droplets are of particular interest as models for minimal synthetic cells. Several studies have demonstrated that catalysis is enhanced or modulated within droplets, but often without turnover of the droplet material itself [5–11]. Importantly, progress has been made in programming dynamics of the droplets themselves; for instance, fuel-driven reaction cycles have been used to control growth and decay of peptide-RNA droplets [12] and such reaction cycles could in turn modulate DNAzyme activity [13]. Theoretical work shows that active droplets should be capable of sustained growth and division under appropriate conditions [14].

However, a critical limitation persists. In these active droplets, dynamic behaviors are typically controlled by fuels and enzymes, rather than by information encoded within the droplet’s own components. What is still missing is thus a direct link between heritable genetic information and the macroscopic behavior of the compartment, a prerequisite for creating active droplets that could eventually evolve. An evolvable system, in turn, should have the capacity to adapt and show emergent behavior, a milestone towards engineering life [2]. RNA is the ideal candidate to realize genetically encoded evolvable and active droplets: On the one hand, RNA can function as an information carrier and it can catalyze chemical reactions. Ribozymes, catalytically active RNA molecules, have been shown to perform ligation, cleavage, and even primitive replication and peptide bond formation [15–21]. They have already been used to control the state of RNA droplets [11]. On the other hand, recent advances in RNA nanotechnology allow for the design of sequence-encoded droplets [22]. It is important to note that most condensates formed from nucleic acids are driven by nonspecific electrostatic interactions between oppositely charged macromolecules, meaning there is typically little sequence specificity in these systems [23–25]. RNA nanotechnology, however, allows for the design of highly specific “nanostar” architectures that interact with one another based on sequence-complementary interactions between terminal loops, the so-called kissing loops, in absence of a polycation [26–28], hence increasing the information that evolution could select for. Such RNA droplets assemble during transcription from a DNA template and have already been equipped with aptamers for sequence-specific recruitment of a substrate to the droplet [26, 27]. Recently, transcription of RNA droplets has been achieved in bacterial cells [29]. We have shown that RNA droplets encapsulate their own DNA template, effectively establishing a link between genotype and phenotype [30]. Moreover, recent works have achieved the dissolution of RNA droplets by strand displacement [28, 31].

To date, however, such genetically-encoded RNA droplets act as passive scaffolds, no active droplets, with chemical turnover, have been realized based on sequence-specific interactions instead of charge multivalency. In this work, we fuse the informational and catalytic properties of RNA to create transiently active, genetically programmed droplets. We achieve this by embedding ribozyme-mediated turnover directly into the architecture of the compartment-forming RNA nanostars. By incorporating specific cleavage sites into the nanostar design, we demonstrate programmable droplet dissolution triggered by trans-acting ribozymes. To study individual droplets over longer periods of time and to prevent fusion events whilst flushing we trap RNA droplets in hydrogel cages. We show that the dissolution kinetics can be tuned by selecting different ribozymes and that we can regrow droplets from recycled DNA templates upon cleavage, effectively establishing a simple dissolution and regrowth cycle. Furthermore, by targeting a chimeric linker molecule in a mixed-droplet population, we use sequence-specific cleavage to induce the segregation of orthogonal droplets. These findings establish a direct, genetically encoded mechanism for controlling the stability, composition, and organization of RNA droplets, representing a key step toward the engineering of responsive and evolvable materials for synthetic biology and protocell research.

## Results and Discussion

To implement genetically encoded ribozyme-regulated droplet dynamics, we built on previously established RNA droplets that form from self-assembling four-arm RNA nanostars [26]. The RNA nanostars are transcribed from a 265 basepair long double-stranded DNA (dsDNA) template using T7 RNA polymerase. They fold co-transcriptionally, i.e. during transcription, into the designed monomeric four-arm shape. Since each arm is equipped with a terminal self-complementary kissing-loop (KL), droplets form by sequence-specific interactions between monomers (Figure 1a). The incorporation of a fluorescent light-up aptamer (FLAP), namely a malachite green aptamer (MGA), enables fluorescence visualization of the droplet behavior. A key advantage is that transcription, folding, and droplet formation occur in a single one-pot reaction at 37 ^°^C. No additional steps, such as purification, component addition or temperature adjustments, are needed. Moreover, the droplet formation is sequence-encoded in the DNA template and not triggered by electrostatic interactions between oppositely charged polymers, which is key towards our aim to link genetic information to droplet behavior.

**Figure 1.**
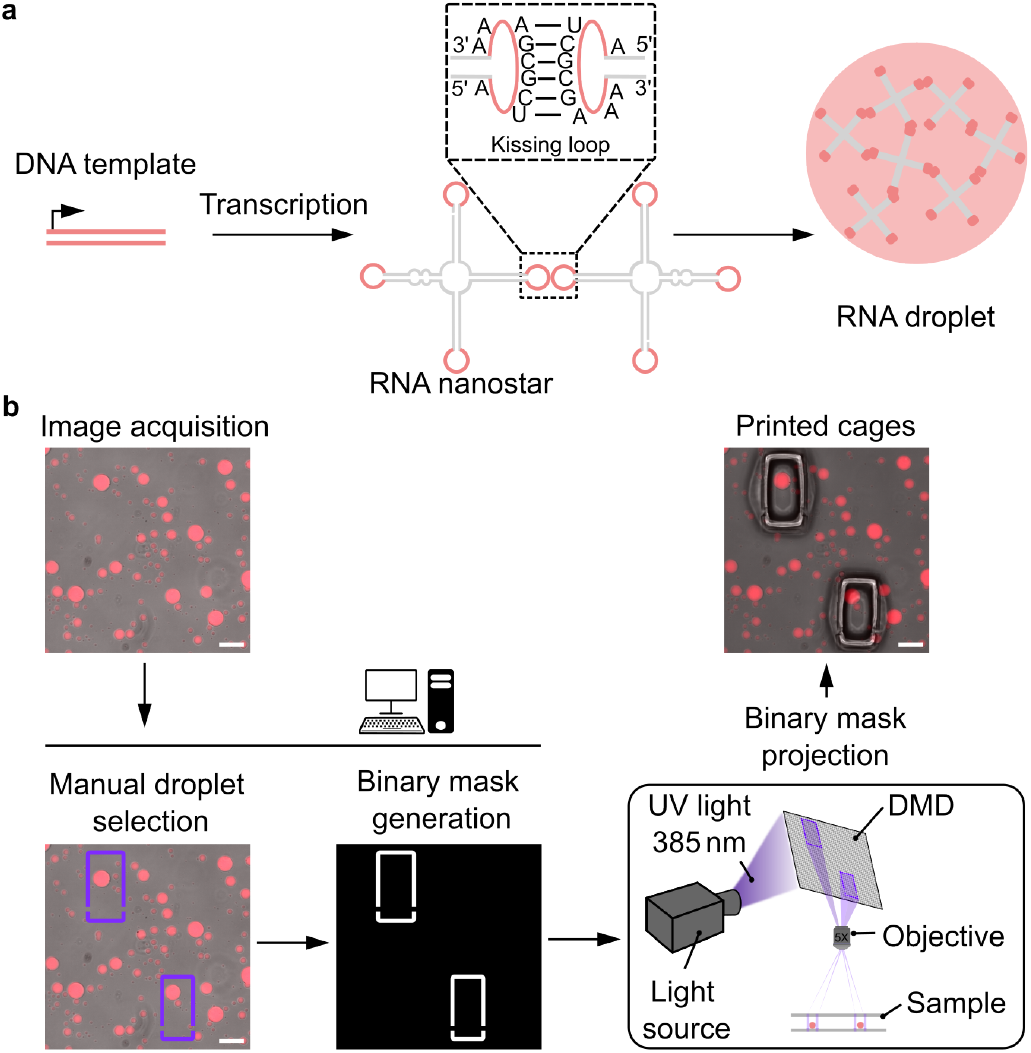
Design and *in situ* trapping of RNA droplets. **a)** Sketch of RNA droplet design and formation. A DNA template codes for a co-transciptionally folding four-arm RNA nanostar, which forms a droplet due to self-complementary kissing loop interactions (dashed box) [26]. **b)** Custom-built hardware and software environment for droplet trapping. A custom-written software interface guides the user through image acquisition and droplet selection. A binary mask is automatically generated and projected onto the sample via a digital micromirror device (DMD) to induce hydrogel cage formation around the selected droplets by targeted photopolymerization. Fluorescent micrographs (fluorescence and brightfield overlay) of RNA droplets illustrate the experimental pipeline (droplets contain MGA and the malachite green dye, *λ*_*ex*_ = 640 nm). Scale bars: 100 µm.

### In situ trapping of RNA droplets

In order to study droplet dynamics in a quantitative manner, we require a method to observe individual droplets over extended periods of time that allows for buffer exchange. Previously, active droplets were trapped in waterin-oil droplets for monitoring purposes[32]. However, in this system, the surrounding oil-phase makes perfusion steps for addition of reagents challenging to realize. In the field of microfabrication, digital micromirror devices (DMDs) have been used to create custom hydrogel microstructures by photopolymerization in order to filter and cage cells [33, 34].

We thus repurposed a semi-automated platform, developed in house [35, 36], for *in situ* trapping of individual RNA droplets inside of hydrogel chambers by targeted photopolymerization. For this purpose, we synthesized a sugar-based photoresist consisting of the monomer glycidyl methacrylate derivatized dextran and the photoinitiator lithium phenyl-2,4,6-trimethylbenzoylphosphinate. We confirmed that RNA droplets are transcribed as before in the photoresist-containing solution (Figure 1b, top left). To allow for targeted photopolymerization around individual droplets, we coupled a DMD into the beam path of a fluorescence microscope as illustrated in Figure 1b. We developed a software interface where users can select individual droplets in a live brightfield or fluorescence image with a simple mouse-click. The rectangular cages are translated into a binary mask, which defines the illumination pattern of the DMD to locally activate the photoinitiator. We printed rectangular cages with manually adjusted dimensions around individual droplets, providing the droplets with sufficient space for growth and dissolution (Figure 1b right). The binary masks defining the rectangular cages include small side inlets that permit reagents to flow inside and reach the trapped droplets. This design allows different reagents to be introduced sequentially into the droplet-containing cages. The droplets remain securely inside the cages and can be monitored for several hours, unlike droplets outside the cages which fuse and drift out of the field of view, in particular when flushing reagents (Figure S1 and Video S1).

### Ribozyme-based RNA droplet dissolution

Having established a reliable setup to trap individual droplets and monitor them over time, we can observe droplet dynamics. Our goal was to introduce droplet dissolution by ribozyme activity. We thus incorporated two ribozyme cleavage sites into the RNA nanostar design to cleave it into two parts, each containing two arms. In particular, we chose cleavage sites for a trans-cleaving ribozyme, which allows us to trigger cleavage upon addition of the ribozyme. Cleavage of the RNA nanostar leads to droplet dissolution because the resulting two-arm nanostars cannot maintain the network (Figure 2a and b).

**Figure 2.**
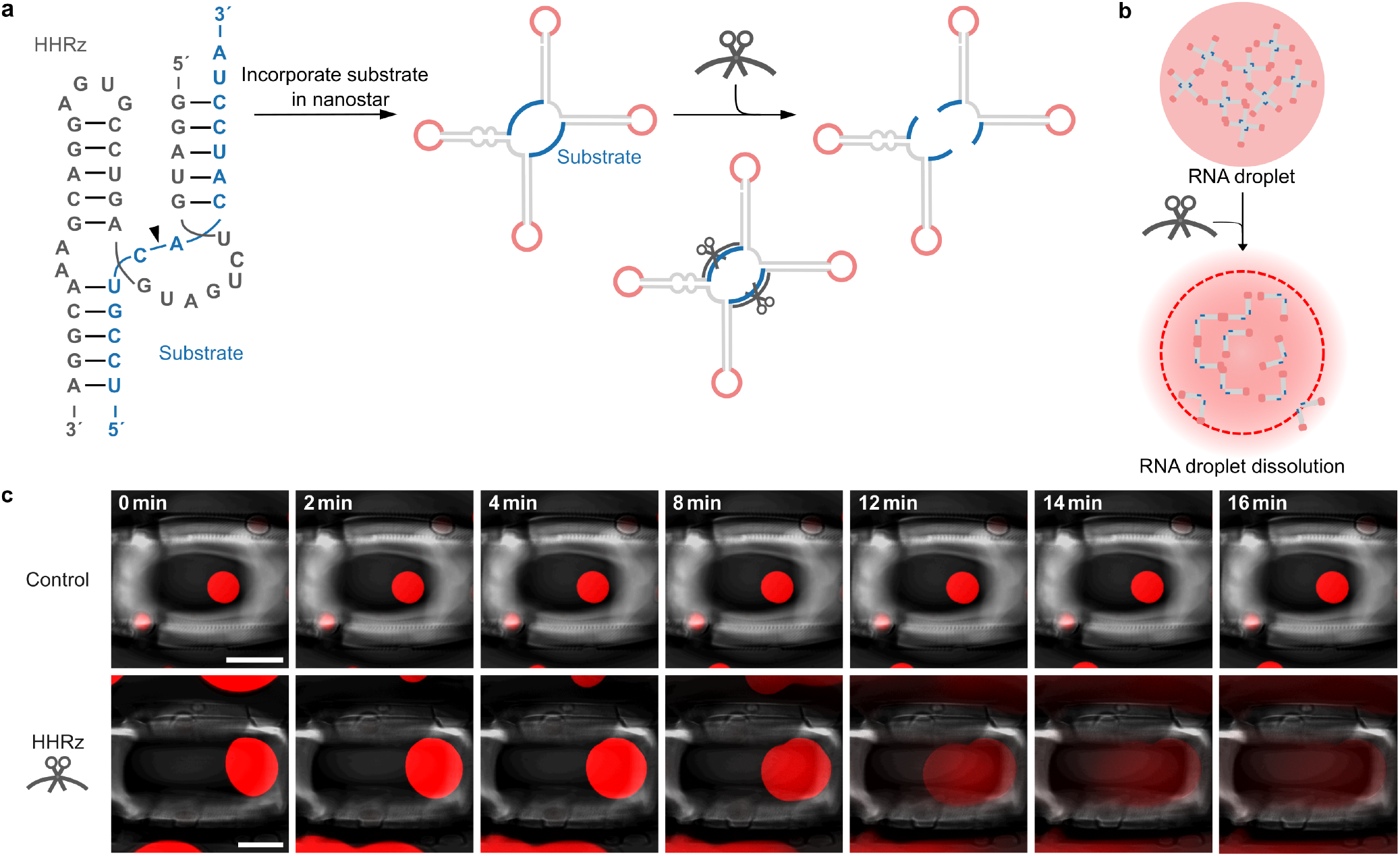
Ribozyme-triggered RNA droplet dissolution. **a)** RNA sequence of the HHRz (gray) and substrate (blue). The cleavage site is indicated with a black arrow. Schematic representation illustrating the incorporation of the substrate sequence into the nanostar design. Upon addition of the ribozyme, cleavage occurs, resulting in two separate fragments, each containing two arms. **b)** Schematic of droplet-forming four-arm RNA nanostars with the incorporated ribozyme cleavage site. Cleaving the monomers into lower-valency (two-arm) structures upon ribozyme addition leads to droplet dissolution. **c)** Confocal timeseries of caged RNA droplets (fluorescence and brightfield overlay). Droplets contain MGA and malachite green dye, *λ*_*ex*_ = 640 nm. At *t* = 0, IVT-buffer (control, top row) or HHRz (bottom row) was added. Scale bars: 50 µm.

We selected a minimal version of the hammerhead ribozyme (HHRz) (Figure 2a), with a trans-Hoogsteen base-pairing interaction, which has been reported to enhance catalytic activity [37]. We chose this version to reduce the sequence length that needs to be incorporated into the nanostar structure, while maintaining the catalytic efficiency of the full-length HHRz. To ensure efficient cleavage, we redesigned the RNA nanostar sequence, such that the ribozyme substrate sequence remains unpaired in the nanostar structure.

The RNA nanostar with the incorporated ribozyme target site (DrA_HHRz_) was transcribed at 37 ^°^C for 24 h. The resulting droplets appeared similar to those without a target site, indicating that the incorporation of a longer single-stranded region (15 nt), does not interfere with droplet formation (Figure 2c). We then confirmed the cleavage of DrA_HHRz_ by the ribozyme via denaturing polyacrylamide gel electrophoresis (PAGE) (Figure S2).

To monitor droplet dissolution, individual droplets were first caged, and then the ribozyme was flushed into the sample. All observed droplets (n = 7) began to dissolve within 15 minutes after the addition of the ribozyme (Figure 2c, Figure S3 and Video S2). This fast dissolution is consistent with the reported cleavage kinetics of the HHRz[37]. *In situ* transcription of the HHRz along side already-formed RNA droplets also resulted in rapid droplet dissolution (Figure S3b).

Whilst for the negative control, where we added IVT-buffer (see Supporting Information), droplets were stable over time, proving that the observed dissolution is indeed an effect of ribozyme cleavage and not due to buffer addition (Figure 2c).

To explore whether this effect can also be observed at the macroscopic scale, we transcribed the RNA droplets in a PCR tube resulting in the formation of a visible gel at the bottom. Addition of the ribozyme led to complete dissolution of the gel (Figure S4).

### Tuning ribozyme activity to control droplet behavior

Next, we set out to tune the ribozyme kinetics. To slow down droplet dissolution to allow for more detailed monitoring, we decided to use a hairpin ribozyme (HPRz), which is reported to have a slower cleavage rate compared to the HHRz [38]. The structure of the HPRz consists of four helical elements (H1 to H4) and two internal loops (A and B) [39]. These two independent folding domains must interact for the ribozyme-substrate complex to be catalytically active. Additionally, the HPRz exhibits biphasic kinetics, with a slow phase arising from reversible substrate binding to the inactive complex [40].

First, cleavage of DrA_HPRz_ by the ribozyme was again confirmed using PAGE (Figure S5). Again, to monitor individual droplets over time, they were trapped in a hydrogel cage, and one of three solutions was added: IVT-buffer (control), HPRz or the IVT containing the DNA template for the HPRz, transcribing the ribozyme *in situ*. Upon addition of the HPRz, the droplets began to shrink gradually over several hours, occasionally accompanied by transient vacuole formation as visible in the confocal timeseries (Figure 3b middle, Figure S6b middle and Video S3) and when quantifying the vacuole-to-area ratio (Figure S7). As expected, dissolution is considerably slower compared to the results for the addition of the HHRz (Figure 2).

**Figure 3.**
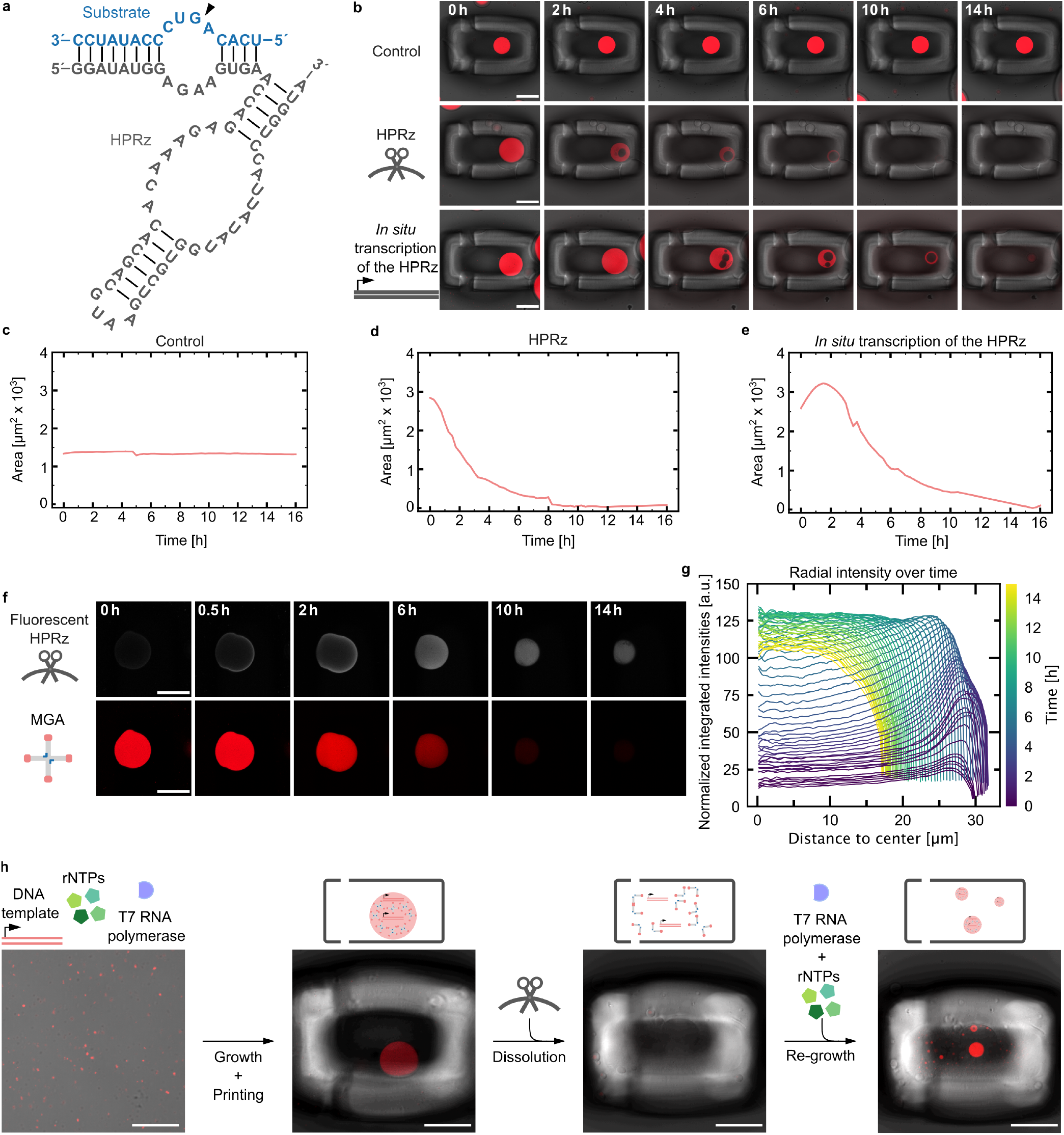
Sequence-encoded control of ribozyme kinetics. **a)** RNA sequence of the HPRz (gray) and substrate (blue). The cleavage site is indicated with a black arrow. **b)** Confocal timeseries of caged RNA droplets (fluorescence and brightfield overlay). At *t* = 0 IVT-buffer (top row), HPRz (middle row) or *in situ* transcription of the HPRz (bottom row) was added. **c-e)** Droplet area for the droplets in b) over time. **f)** Confocal timeseries of caged RNA droplets after the addition of fluorescently labeled HPRz. At *t* = 0, the HPRz was flushed in. The HPRz (white) is labeled with fluorescein, *λ*_*ex*_ = 488 nm. **g)** Radial intensity profiles showing the diffusion of fluorescently labeled HPRz into a single RNA droplet over time (see Supporting Information). Normalized integrated intensities were measured as a function of distance from the droplet center. The color gradient represents different time points over a 14 h period. Additional replicates are shown in Figure S9. **h)** Schematic representation and confocal micrographs (fluorescence and brightfield overlay) showing a minimal cycle of growth, dissolution and regrowth. Key steps include transcription and droplet formation, HPRz-induced dissolution, and droplet regrowth. For all confocal micrographs droplets (red) contain MGA and malachite green dye, *λ*_*ex*_ = 640 nm. Scale bars: 50 µm.

Interestingly, when transcribing the ribozyme *in situ*, the droplets initially grew before shrinkage set in (Figure 3b lower, Figure S6a lower and Video S4), suggesting that more RNA nanostars were also transcribed. This behavior aligns with our previous findings [30], which showed that the DNA template for the RNA nanostar is encapsulated within the droplets. Consequently, when the ribozyme is transcribed *in situ*, additional new RNA droplet material is transcribed along with it. Because the HPRz cleaves more slowly, this new production of RNA competes with cleavage, leading to an initial increase in droplet size before ribozyme concentration increases and thus dissolution takes over (Figure 3c-e and Figure S6-S8).

To investigate the diffusion dynamics and mode of action of the ribozyme within RNA droplets, we labeled the ribozyme by integration of fluorescent UTPs (see Supporting Information) and monitored its spatial distribution using confocal microscopy (Figure 3f). Initially, the ribozyme accumulates at the droplet periphery, then gradually diffuses inward over several hours, coinciding with droplet dissolution (Figure 3f and Figure S9). We quantified this behavior by measuring radial fluorescence intensity profiles over time. The signal initially peaks at the periphery and progressively shifts toward the center of the droplet as the droplet shrinks (Figure 3g and Figure S9). This inward propagation suggests that cleavage activity is initially localized near the droplet surface, constrained by the internal architecture, and continues throughout the droplet as dissolution progresses.

We further tested whether RNA droplets, once dissolved, could regrow upon addition of fresh IVT-buffer and T7 RNA polymerase, but without adding new DNA template. Indeed, new droplets formed, suggesting that the DNA released during cleavage can be recycled for transcription and that DNA-encoded growth-dissolution cycles should in principle be possible (Figure 3h). We also verified the sequence specificity of ribozyme cleavage by adding the HHRz to DrA_HPRz_. As expected, the droplets remained intact (Figure S10), confirming the ribozymes’ specificity for their substrate sequence rather than any single-stranded regions. This allows for orthogonal cleavage and further demonstrates that the susceptibility to dissolution can be genetically encoded in the DNA template.

### RNA droplet segregation by ribozyme activity

Multiple RNA droplet species can be simultaneously transcribed using orthogonal KL sequences, and distinguished by confocal microscopy through the incorporation of distinct FLAPs in their nanostar designs [26, 27]. Here we use two other nanostar designs from *Fabrini et al*. [26]. The first one, DrB was reported to form droplets orthogonal to DrA and is functionalized with the Broccoli aptamer (BrA) [26]. The second is a chimeric linker RNA (L) containing two KLs for DrA and two KLs for DrB. When included in a DNA template ratio of 1:2:1 (DrA: L: DrB), the system produces homogeneously mixed droplets (Figure S11).

We hypothesized that we could achieve segregation of the homogeneous droplet by cleaving the linker with our previously established ribozyme cleavage strategy. We thus incorporated the substrate sequence for the highly efficient HHRz into L (L_HHRz_) following the same design strategy as for DrA_HHRz_. The ribozyme cleavage site was positioned in such a way that the resulting fragments either contain KLs A or KLs B. Upon addition of the HHRz, the linker should be cleaved specifically, while DrA and DrB nanostars remain intact. Complete linker cleavage should disrupt DrA-DrB interactions, leading to segregation into two distinct droplets (Figure 4a). We again confirmed cleavage of L_HHRz_ using PAGE (Figure S2).

**Figure 4.**
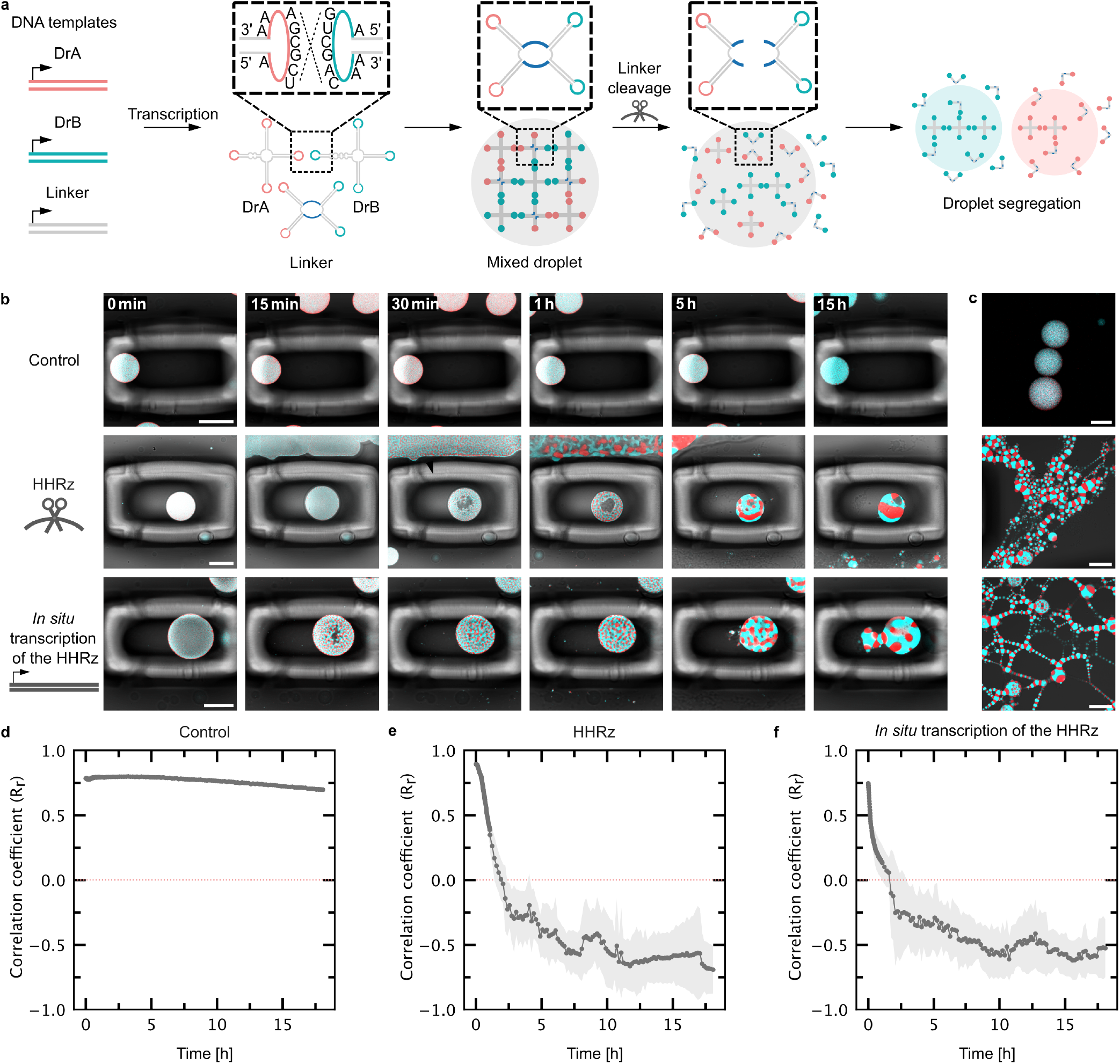
RNA droplet segregation with HHRz. **a)** Schematic of the cleavage strategy. Droplets are formed from two orthogonal RNA nanostars DrA and DrB with a chimeric linker (L) containing two KLs of DrA and two KLs of DrB[26]). We modified L with a HHRz cleavage site. Once the HHRz is added, the linker with incorporated substrate sequence is cleaved, leading to droplet segregation. **b)** Confocal timeseries of segregating RNA droplets (fluorescence and brightfield overlay). At *t* = 0, IVT-buffer (top row), over night transcribed ribozyme (middle row) or IVT containing the DNA template for the HHRz, transcribing the ribozyme *in situ* (bottom row) was added. **c)** Confocal micrographs of RNA droplets in uncaged regions acquired after ending the timelapse shown in b). Red droplets contain MGA and malachite green dye, *λ*_*ex*_ = 640 nm, blue droplets contain BrA and DFHBI-1T, *λ*_*ex*_ = 488 nm. Scale bars: 50 µm. **d-f)** Colocalization analysis. Pearson correlation R_r_ values are plotted over time (mean *±* s.d., n = 5 regions).

DrA, L_HHRz_ and DrB were transcribed using a 1:2:1 DNA template ratio. We confirmed that the incorporation of the single-stranded substrate region (15 nt) does not hinder its mixing properties (Figure 4b). Again, to monitor individual droplets over time, they were trapped in photopolymerized cages, and IVT-buffer (control), ribozyme or the IVT containing the DNA template for the HPRz, transcribing the ribozyme *in situ* was added. RNA droplet segregation could be observed 15 to 30 minutes after the addition of the HHRz. Over the course of several hours the droplets first collapsed, accompanied by vacuole formation in the center of the droplet (Figure 4b and Video S5). Complete separation into two droplets was not observed for the monitored droplets, they remained connected at the interface. However, smaller droplets that were not in focus in the caged region, as well as uncaged droplets, were fully segregated, forming long chains (Figure 4c and Figure S12). This suggests that shear forces from flow in the microfluidic chamber may facilitate segregation outside the caged regions. From experiments with the HHRz dissolving DrA we know that the cleavage reaction is highly efficient and occurs within minutes. We therefore assumed that cleavage of the linker was highly efficient and attributed the incomplete segregation of the RNA droplets to the viscosity of the droplets which slows down the separation process, and non-specific interactions between the monomers. To quantitatively describe the segregation kinetics, we determined the Pearson’s correlation coefficient (R_r_) over time as a measure of colocalization between DrA and DrB. A value of R_r_ = 1 corresponds to perfect colocalization, whereas R_r_= − 1 represents perfectly anticorrelated data. Initially, R_r_ decreased rapidly within the first two hours before stabilizing around − 0.5. Together, these results confirm that ribozyme-mediated linker cleavage effectively triggers droplet segregation. We wanted to further investigate the efficiency of the ribozyme approach by comparing it to enzymatic segregation strategies using RNase H. As RNase H is known to specifically degrade the RNA in basepaired DNA-RNA hybrids, we added RNase H together with a DNA strand complementary to the single stranded RNA region in L_HPRz_ [41, 42]. As expected, RNase H cleaved the linker, leading to demixing of the RNA droplets. Remarkably, we found that the catalytically active ribozyme performed just as efficiently as RNase H in cleaving the linker and disrupting droplet interactions (Figure S13). Again, these observations suggest that un-cleaved linker is unlikely to be the cause for incomplete segregation of the droplets. Notably, the ribozyme-cleavage strategy thus allowed us to genetically encode the droplet’s capacity to segregate, whereas RNase H would degrade any DNA-RNA hybrid. To confirm that segregation is not due to non-specific cleavage of the single-stranded region of the linker, we studied the stability of the mixed droplets over time. Mixed droplets (using L_HPRz_) remained stable for several days, with non-specific segregation occurring only after 6 days, likely due to degradation of the single-stranded RNA regions in L (Figure S14). Since our experiments focus on effects within a 48-hour window, we conclude that spontaneous segregation is negligible in this time frame.

## Conclusion

In this study, we demonstrate that ribozyme catalysis can serve as a genetically encoded mechanism to control the stability, composition, and organization of RNA-based droplets. To study the dissolution kinetics of RNA droplets, we repurposed an in-house developed semi-automated platform [35, 36] to trap individual droplets *in situ* through local photopolymerization. By embedding ribozyme target sequences into self-assembling RNA nanostars, we achieved droplet dissolution and drove the segregation of mixed droplet populations. Crucially, we were able to modulate the dissolution kinetics by selecting different ribozymes and their respective target sequences. We find that the HHRz drives rapid droplet disassembly within minutes, while the HPRz acts on timescales of hours. When transcribed in situ, the HPRz exhibits a dynamic interplay between droplet growth and degradation, suggesting the potential for designing self-regulating systems in which assembly, disassembly, and reassembly occur in a cyclic fashion. Achieving such sustained, life-like cycles of growth and division will require further development of mechanisms to independently tune the rates of these competing processes.

As the field of active droplets advances, we envision that the droplet trapping technology by photopolymerization will provide a powerful tool for synthetic cell research, enabling precise monitoring and manipulation of synthetic cells, with minimal perturbation for longer periods of time.

Because RNA droplets are directly transcribed from DNA templates, the information encoded in their sequence directly determines their macroscopic behavior. Due to the sequence specificity of ribozyme cleavage, only droplets containing the matching substrate are subject to dissolution, while mutated droplets should escape catalysis. Note that droplet segregation has previously been achieved for DNA droplets, triggered either by light [42] or RNase H activity [41, 42]. In these cases, however, segregation was not linked to a specific sequence. Linking DNA-encoded information to the droplet’s capacity to segregate opens a path toward genotype-dependent selection and evolution in synthetic RNA compartments.

We further show that RNA droplets, once dissolved, can regrow without the addition of new DNA template, suggesting that the DNA released upon cleavage can be recycled for droplet formation. This establishes a minimal cycle of growth and decay, encoded entirely in nucleic acid sequences. Beyond controlling individual droplets, we demonstrate that ribozyme activity can induce transitions between mixed and demixed states. Such programmable segregation lays the foundation for engineering RNA droplets capable of asymmetric division. Just as living cells employ tightly regulated mechanisms for division, ribozyme-mediated cleavage offers a genetically encoded strategy to split condensates into distinct, self-contained compartments that retain the capacity for regrowth.

A defining feature of our system is that its dynamic behavior does not rely on externally supplied fuel consumption. Instead, disassembly is an intrinsic property encoded within the RNA sequence, with the energy for turnover released via ribozyme-catalyzed phosphodiester bond cleavage. In this sense, our droplets already represent a genetically programmed form of transiently active condensates, linking heritable genetic information and catalytic activity, thus laying the foundation for evolving catalytically active droplets. At the same time the repertoire of ribozymes goes beyond phosphodiester bond cleavage as ribozymes are also capable of fuel-consuming reactions using cofactors like S-adenosyl methionine (SAM) [43]. Harnessing such reactions could couple droplet turnover to fuel consumption, providing a route toward more complex RNA-based compartments with integrated metabolic activity. Looking further forward, the versatility of ribozymes offers numerous possibilities. Beyond cleavage, ribozymes capable of ligation or polymerization could be incorporated to support self-sustained RNA synthesis [8], advancing toward synthetic protocells driven entirely by RNA catalysis. Future *in vitro* selection of ribozymes could focus on their catalytic activity within the highly charged environment of condensates, rather than in bulk. In this way, our work provides a genetically encoded route towards fuel-dependent active droplets, closing the gap between passive RNA condensates and protocell-like systems with evolvable, life-like behavior.

## Acknowledgements

The authors thank Mai Phuong Tran and Cody Geary for helpful discussions related to this work. This work was supported by the ERC Starting Grant “ENSYNC” (No. 101076997), Deutsche Forschungsgemeinschaft (DFG, German Research Foundation) under Germany’s Excellence Strategy via the Excellence Cluster 3D Matter Made to Order (EXC-2082/1 – 390761711) and under CRC 392 as well as a Research Grant from HFSP (Ref.-No: RGP003/2023,

DOI:https://doi.org/10.52044/HFSP.RGP0032023.pc.gr.168589). The authors thank the Alfried Krupp von Bohlen und Halbach Foundation for generous support. T.A. and F.G. thank the Carl Zeiss Foundation for financial support.

## References

1. Poudyal, R. R., Pir Cakmak, F., Keating, C. D. & Bevilacqua, P. C. Physical Principles and Extant Biology Reveal Roles for RNA-Containing Membraneless Compartments in Origins of Life Chemistry. Biochemistry 57.PMID: 29560725, 2509–2519. eprint: 10.1021/acs.biochem.8b00081. https://doi.org/10.1021/acs.biochem.8b00081(2018).

2. Kriebisch, C. M. E. et al. A roadmap toward the synthesis of life. English. Chem 11.Publisher: Elsevier. issn: 2451-9294, 2451-9308. https://www.cell.com/chem/abstract/S2451-9294(24)00644-2 (2025) (Mar. 2025).

3. Hyman, A. A., Weber, C. A. & Jülicher, F. Liquid-liquid phase separation in biology. Annual review of cell and developmental biology 30,39–58 (2014).

4. Beneyton, T. et al. Out-of-equilibrium microcompartments for the bottom-up integration of metabolic functions. Nature communications 9,2391 (2018).

5. Drobot, B. et al. Compartmentalised RNA catalysis in membrane-free coacervate protocells. Nature communications 9,3643 (2018).

6. Poudyal, R. R., Keating, C. D. & Bevilacqua, P. C. Polyanion-assisted ribozyme catalysis inside complex coacervates. ACS chemical biology 14,1243–1248 (2019).

7. Poudyal, R. R. et al. Template-directed RNA polymerization and enhanced ribozyme catalysis inside membraneless compartments formed by coacervates. Nature communications 10,490 (2019).

8. Le Vay, K., Song, E. Y., Ghosh, B., Tang, T.-Y.D. & Mutschler, H. Enhanced ribozyme-catalyzed recombination and oligonucleotide assembly in peptide-RNA condensates. Angewandte Chemie International Edition 60,26096–26104 (2021).

9. Iglesias-Artola, J. M. et al. Charge-density reduction promotes ribozyme activity in RNA–peptide coacervates via RNA fluidization and magnesium partitioning. Nature chemistry 14,407–416 (2022).

10. Küffner, A. M. et al. Acceleration of an enzymatic reaction in liquid phase separated compartments based on intrinsically disordered protein domains. ChemSystemsChem 2,e2000001 (2020).

11. Le Vay, K. K., Salibi, E., Ghosh, B., Tang, T. D. & Mutschler, H. Ribozyme activity modulates the physical properties of RNA–peptide coacervates. Elife 12,e83543 (2023).

12. Donau, C. et al. Active coacervate droplets as a model for membraneless organelles and protocells. Nature communications 11,5167 (2020).

13. Holtmannspötter, A.-L. et al. Regulating Nucleic Acid Catalysis Using Active Droplets. Angewandte Chemie International Edition 63,e202412534 (2024).

14. Zwicker, D., Seyboldt, R., Weber, C. A., Hyman, A. A. & Jülicher, F. Growth and division of active droplets provides a model for protocells. en. Nature Physics 13.Publisher: Nature Publishing Group, 408– 413. issn: 1745-2481. https://www.nature.com/articles/nphys3984) (2025 (Apr. 2017).

15. Kruger, K. et al. Self-splicing RNA: autoexcision and autocyclization of the ribosomal RNA intervening sequence of Tetrahymena. cell 31,147–157 (1982).

16. Guerrier-Takada, C., Gardiner, K., Marsh, T., Pace, N. & Altman, S. The RNA moiety of ribonuclease P is the catalytic subunit of the enzyme. Cell 35,849–857 (1983).

17. Johnston, W. K., Unrau, P. J., Lawrence, M. S., Glasner, M. E. & Bartel, D. P. RNA-catalyzed RNA polymerization: accurate and general RNA-templated primer extension. Science 292,1319–1325 (2001).

18. Horning, D. P. & Joyce, G. F. Amplification of RNA by an RNA polymerase ribozyme. Proceedings of the National Academy of Sciences 113,9786–9791 (2016).

19. Zhang, B. & Cech, T. R. Peptide bond formation by in vitro selected ribozymes. Nature 390,96–100 (1997).

20. Zhou, L., O’Flaherty, D. K. & Szostak, J. W. Assembly of a ribozyme ligase from short oligomers by nonenzymatic ligation. Journal of the American Chemical Society 142,15961–15965 (2020).

21. Nomura, Y. & Yokobayashi, Y. RNA ligase ribozymes with a small catalytic core. Scientific Reports 13,8584 (2023).

22. Monari, L., Braun, I., Poppleton, E. & Göpfrich, K. PyFuRNAce: An integrated design engine for RNA origami. bioRxiv. eprint: https://www.biorxiv.org/content/early/2025/04/18/2025.04.17.647389.full.pdf. https://www.biorxiv.org/content/early/2025/04/18/2025.04.17.647389 (2025).

23. Majumder, S., Coupe, S., Fakhri, N. & Jain, A. Sequence-encoded intermolecular base pairing modulates fluidity in DNA and RNA condensates. en. Nature Communications 16.Publisher: Nature Publishing Group, 4258. issn: 2041-1723. https://www.nature.com/articles/s41467-025-59456-0 (2025) (May 2025).

24. Nguyen, H. T., Hori, N. & Thirumalai, D. Condensates in RNA repeat sequences are heterogeneously organized and exhibit reptation dynamics. en. Nature Chemistry 14.Publisher: Nature Publishing Group, 775–785. issn: 1755-4349. https://www.nature.com/articles/s41557-022-00934-z (2025) (July 2022).

25. Wollny, D. et al. Characterization of RNA content in individual phase-separated coacervate microdroplets. en. Nature Communications 13.Publisher: Nature Publishing Group, 2626. issn: 2041-1723. https://www.nature.com/articles/s41467-022-30158-1 (2025) (May 2022).

26. Fabrini, G. et al. Co-transcriptional production of programmable RNA condensates and synthetic organelles. Nature Nanotechnology, 1–9 (2024).

27. Stewart, J. M. et al. Modular RNA motifs for orthogonal phase separated compartments. Nature Communications 15,6244 (2024).

28. Udono, H. et al. Programmable computational RNA droplets assembled via kissing-loop interaction. ACS nano 18,15477–15486 (2024).

29. Ng, B. et al. Expression of nano-engineered RNA organelles in bacteria en. ISSN: 2692-8205 Pages: 2025.07.08.663582 Section: New Results. July 2025. https://www.biorxiv.org/content/10.1101/2025.07.08.663582v1 (2025).

30. Verstraeten, W. et al. Genetic encoding and mutagenesis of RNA droplet phenotypes en. May 2025. https://chemrxiv.org/engage/chemrxiv/article-details/680ca3e6927d1c2e66de8256(2025).

31. Tang, A. A., Gobry, M. V., Li, S., Andersen, E. S. & Franco, E. Switchable RNA motifs for dynamic transcriptional control of RNA condensates. Nucleic Acids Research 53,gkaf497. issn: 1362-4962. 10.1093/nar/gkaf497 (2025) (July 2025).

32. Bergmann, A. M. et al. Evolution and Single-Droplet Analysis of Fuel-Driven Compartments by Droplet-Based Microfluidics. Angewandte Chemie International Edition 61,e202203928. eprint: https://onlinelibrary.wiley.com/doi/pdf/10.1002/anie.202203928. https://onlinelibrary.wiley.com/doi/abs/10.1002/anie.202203928 (2022).

33. Yang, W., Yu, H., Liang, W., Wang, Y. & Liu, L. Rapid Fabrication of Hydrogel Microstructures Using UV-Induced Projection Printing. en. Micromachines 6.Publisher: Multidisciplinary Digital Publishing Institute, 1903–1913. issn: 2072-666X. https://www.mdpi.com/2072-666X/6/12/1464 (2025) (Dec. 2015).

34. Zhang, H. et al. Rapid trapping and tagging of microparticles in controlled flow by in situ digital projection lithography. en. Lab on a Chip 22.Publisher: Royal Society of Chemistry, 1951–1961. https://pubs.rsc.org/en/content/articlelanding/2022/lc/d2lc00186a (2025) (2022).

35. GÖPFRICH, K. et al. en. WO2023052442A1. https://patents.google.com/patent/WO2023052442A1/en?inventor=tobias+abele&oq=tobias+abele (2025)(2023).

36. GÖPfrich, K., Abele, T. & Maurer, S. en. WO2024256656A1. https://patents.google.com/patent/WO2024256656A1/en?inventor=tobias+abele&assignee=MAX-PLANCK-Gesellschaft+zur+F%C3%B6rderung+der+Wissenschaften+e.V. (2025) (2024).

37. O’Rourke, S. M., Estell, W. & Scott, W. G. Minimal hammerhead ribozymes with uncompromised catalytic activity. Journal of molecular biology 427,2340–2347 (2015).

38. Walter, N. G. & Burke, J. M. The hairpin ribozyme: structure, assembly and catalysis. Current opinion in chemical biology 2,303 (1998).

39. Butcher, S. E., Heckman, J. E. & Burke, J. M. Reconstitution of Hairpin Ribozyme Activity following Separation of Functional Domains (). Journal of Biological Chemistry 270,29648–29651 (1995).

40. Esteban, J. A., Banerjee, A. R. & Burke, J. M. Kinetic mechanism of the hairpin ribozyme: identification and characterization of two nonexchangeable conformations. Journal of Biological Chemistry 272,13629– 13639 (1997).

41. Sato, Y., Sakamoto, T. & Takinoue, M. Sequence-based engineering of dynamic functions of micrometer-sized DNA droplets. Science advances 6,eaba3471 (2020).

42. Tran, M. P. et al. A DNA segregation module for synthetic cells. Small 19,2202711 (2023).

43. Scheitl, C. P. M., Ghaem Maghami, M.Lenz, A.-K. & Höbartner, C. Site-specific RNA methylation by a methyltransferase ribozyme. en. Nature 587.Publisher: Nature Publishing Group, 663–667. issn: 1476-4687. https://www.nature.com/articles/s41586-020-2854-z (2025) (Nov. 2020).

