## Supplementary Information for "Growth, dissolution and segregation of genetically encoded RNA droplets by ribozyme catalysis"

**4 Uncropped and non-inverted gel images 26**

### 1 Supplementary Tables

#### 1.1 DNA Sequences (coding strand 5´-3')

DrA:

```
  1 GGGCATTCTA ATACGACTCA CTATAGGACA GTGCTATGAG
 41 TGTGCACGGG ATCCCGACTG GCCGCATCGC GAAAGTGGCC
 81 AGGTAACGAA TGGATCCTGT GCTGCACATT AGAGTCGCTG
121 TATGACCCAT CGCGAAAGGG TCGTACAGCG GCTCTAGTGT
161 GCTGCACAGT GTCTGTGCGA CTGCACGCAT CGCGAAAGCG
201 TGTAGTCGCA TAGACATTGT GCTCACTCGT AGCATTGTCC
241 CTGTCTCCAT CGCGAAAGGA GATAG
```

DrA$_{\mathrm{HHRz}}$:

```
  1 GGGCATTCTA ATACGACTCA CTATAGGACA GTGCTATGAG
 41 TGTTCCGTCA CATCCTATGC ACGGGATCCC GACTGGCCGC
 81 ATCGCGAAAG TGGCCAGGTA ACGAATGGAT CCTGTGCTGC
121 ACATTAGAGT CGCTGTATGA CCCATCGCGA AAGGGTCGTA
161 CAGCGGCTCT AGTGTGCTTC CGTCACATCC TATCGACAGT
201 GTCTGTGCGA CTGCACGCAT CGCGAAAGCG TGTAGTCGCA
241 TAGACATTGT CGTCACTCGT AGCATTGTCC CTGTCTCCAT
281 CGCGAAAGGA GATAG
```

DrA$_{\mathrm{HPRz}}$:

```
  1 GGGCATTCTA ATACGACTCA CTATAGGACA GTGCTATGAG
 41 TGTTCACAGT CCCATATTCG CACGGGATCC CGACTGGCCG
 81 CATCGCGAAA GTGGCCAGGT AACGAATGGA TCCTGTGCTG
121 CACATTAGAG TCGCTGTATG ACCCATCGCG AAAGGGTCGT
161 ACAGCGGCTC TAGTGTGCTC ACAGTCCCAT ATCCTGCACA
201 GTGTCTGTGC GACTGCACGC ATCGCGAAAG CGTGTAGTCG
241 CATAGACATT GTGCTCACTC GTAGCATTGT CCCTGTCTCC
281 ATCGCGAAAG GAGATAG
```

DrB:

```
  1 GGGCATTCTA ATACGACTCA CTATAGGACA GTGCTATGAG
 41 TGTCGCGACG GAGACGGTCG GGTCCAGATA GGCCAGTCGA
 81 CAAGGTCTAT CTGTCGAGTA GAGTGTGGGC TCCGTCGCGT
121 GCACATTAGA GTCGCTGTAT GCCACAGTCG ACAAGTGGCG
161 TACAGCGGCT CTAGTGTGCT GCACAGTGTC TGTGCGACTG
201 CACCCAGTCG ACAAGGGTGT AGTCGCATAG ACATTGTGCT
241 CACTCGTAGC ATTGTCCCTG TCTCCAGTCG ACAAGGAGAT
281 AG
```

L:

```
  1 GGGCATTCTA ATACGACTCA CTATAGGACA GTGCTATGAG
 41 TGTGCACAGT GTCTGTGCGA CTGCACCCAT CGCGAAAGGG
 81 TGTAGTCGCA TAGACATTGT GCTGCACATT AGAGTCGCTG
121 TATGAGGGAT CGCGAAACCC TCGTACAGCG GCTCTAGTGT
161 GCTCGCGTGC CTCAGAGGAC CTGTCACCAG TCGACAAGGT
201 GATAGGTCCT TTGAGGTACG CGTCACTCGT AGCATTGTCC
241 CTGTCTCCAG TCGACAAGGA GATAG
```

$L_{HHRz}$:

```
  1 GGGCATTCTA ATACGACTCA CTATAGGACA GTGCTATGAG
 41 TGTTCCGTCA CATCCTATGC ACAGTGTCTG TGCGACTGCA
 81 CCCATCGCGA AAGGGTGTAG TCGCATAGAC ATTGTGCTGC
121 ACATTAGAGT CGCTGTATGA GGGATCGCGA AACCCTCGTA
161 CAGCGGCTCT AGTGTGCTTC CGTCACATCC TATCGCGTGC
201 CTCAGAGGAC CTGTCACGAG TCGACAACGT GATAGGTCCT
241 TTGAGGTACG CGTCACTCGT AGCATTGTCC CTGTCTCCAG
281 TCGACAAGGA GATAG
```

$L_{HPRz}$:

```
  1 GGGCATTCTA ATACGACTCA CTATAGGACA GTGCTATGAG
 41 TGTTCACAGT CCCATATTCT GCACAGTGTC TGTGCGACTG
 81 CACCCATCGC GAAAGGGTGT AGTCGCATAG ACATTGTGCT
121 GCACATTAGA GTCGCTGTAT GAGGGATCGC GAAACCCTCG
161 TACAGCGGCT CTAGTGTGCT TCACAGTCCC ATATCCTCGC
201 GTGCCTCAGA GGACCTGTCA CGAGTCGACA ACGTGATAGG
241 TCCTTTGAGG TACGCGTCAC TCGTAGCATT GTCCCTGTCT
281 CCAGTCGACA AGGAGATAG
```

HHRz:

```
 1 GGGCATTCTA ATACGACTCA CTATAGGATG TCTGATGAGT
41 CCGTGAGGAC GAAACGGA
```

HPRz:

```
 1 GGGCATTCTA ATACGACTCA CTATAGGATA TGGAGAAGTG
41 AACCAGAGAA ACACACGACG TAAGTCGTGG TATATTACCT
81 GGTA
```

DNA$_{c1}$:

```
 1 GAATATGGGA CTGTGA
```

DNA$_{c2}$:

```
 1 GGATATGGGA CTGTGA
```

Table S1: Sequences of the DNA templates. All sequences are written in 5'-3' direction.

#### 1.2 Primer Sequences

| T7 fwd primer: |
|---|
| 1 GGGCATTCTA ATACGACTCA CTATA |

| T7 fwd primer: |
|---|
| 1 GGGCATTCTA ATACGACTCA CTATA |

| DrA rev primer: |
|---|
| 1 CTATCTCCTT TCGCGATGGA |

| DrB rev primer: |
|---|
| 1 CTATCTCCTT GTCGACTGGA G |

| L rev primer: |
|---|
| 1 CTATCTCCTT GTCGACTGGA G |

Table S2: Sequences of the PCR primers. All sequences are written in 5'-3' direction

### 2 Supplementary Figures

#### 2.1 Figure S1: Printed cages prevent droplet fusion and movement

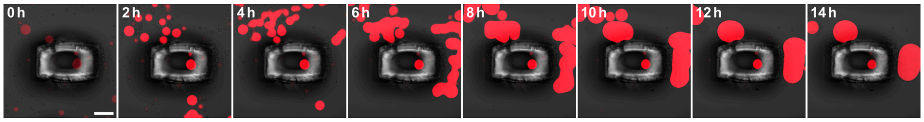

Figure S1: Printed cages prevent droplet fusion and movement. Confocal time-series of caged RNA droplets ($DrA_{HPRz}$, contain MGA and are labeled with malachite green dye, $\lambda_{ex} = 640\,nm$), fluorescence and brightfield overlay), showing that the droplet remains inside of the cage for 14 h. The cage prevents movement and fusion with other droplets and therefore allows to observe individual droplets over time. At $t = 0$, IVT-buffer was flushed in. Scale bars: 50 µm.

#### 2.2 Figure S2: Denaturing polyacrylamide gel electrophoresis of hammerhead ribozyme cleavage products

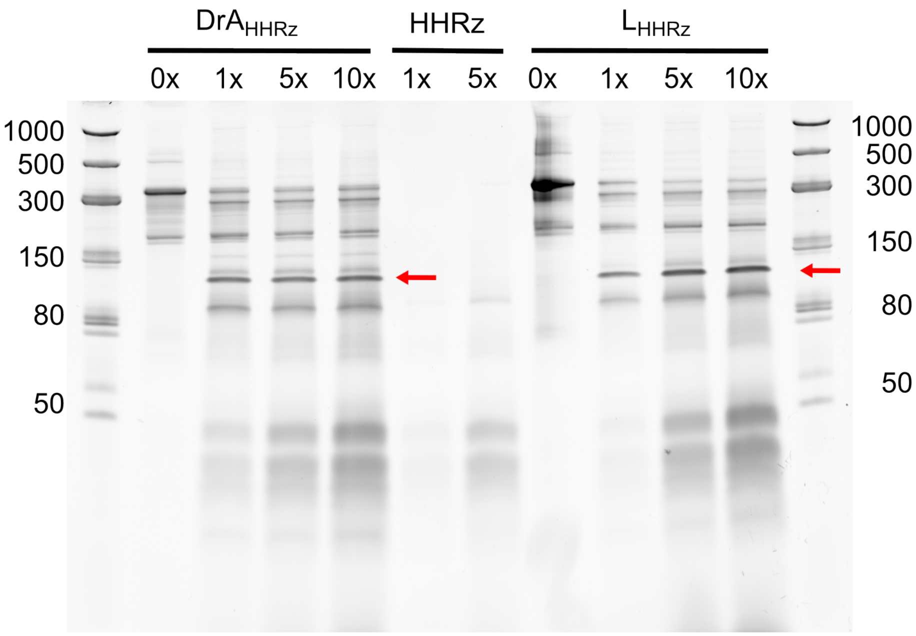

Figure S2: Denaturing polyacrylamide gel electrophoresis of hammerhead ribozyme cleavage products. *In vitro* transcribed RNA nanostars (corresponding design is written on top of the gel) were incubated with increasing volumes of ribozyme (1×, 5×, 10×), where 1× corresponds to 1 µL of ribozyme added to 1 µL of target RNA in a total volume of 50 µL. Lanes labeled 0× serve as cleavage-negative controls with no ribozyme added, the lanes labeled HHRz or HPRz 1× and 5× represent ribozyme-only controls, where ribozymes were diluted (1 µL or 5 µL )in 50 µL IVT-buffer without droplet RNA (for more details see methods). Cleavage products were resolved by 10 % denaturing PAGE stained with GelRed®. Low Range ssRNA Ladder is used as reference, length in bases is reported as text close to the corresponding bands. Red arrows indicate expected cleavage products. Cleavage of full-length constructs (270 nt) is expected to yield three fragments with a lenght of 24 nt, 135 nt and 111 nt.

#### 2.3 Figure S3: Dissolution of RNA droplets induced by a trans-acting hammerhead ribozyme

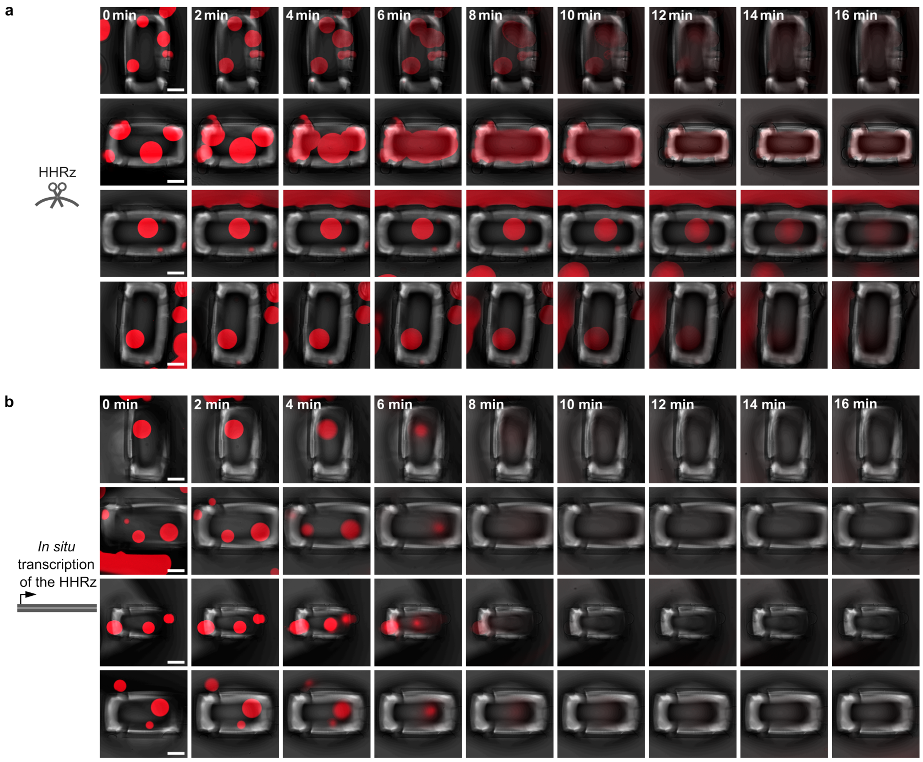

Figure S3: Dissolution of RNA droplets induced by a trans-acting hammerhead ribozyme. Confocal timeseries (fluorescence and brightfield overlay, droplets contain MGA and are labeled with malachite green dye, $\lambda_{ex} = 640\,\text{nm}$) for multiple technical replicates (n=7, 4 are shown here) of dissolving RNA droplets using a trans-acting HHRz. Scale bars: 50 µm. At $t = 0$, the HHRz (**a**) or IVT containing the DNA template for the HHRz, transcribing the ribozyme *in situ* (**b**) was added using a microfluidic pump.

#### 2.4 Figure S4: Dissolution of RNA droplets forming a gel at the bottom of a PCR tube induced by a trans-acting hammerhead ribozyme

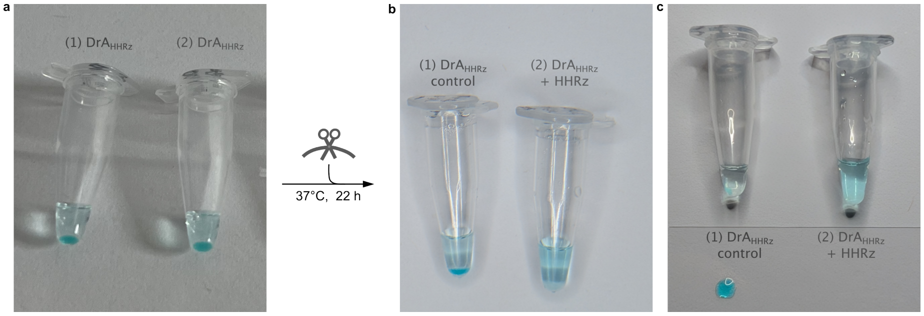

Figure S4: Dissolution of RNA droplets forming a gel at the bottom of a PCR tube induced by a trans-acting hammerhead ribozyme. a) RNA droplets composed of $DrA_{HHRz}$ were formed by IVT in a 20 µL reaction at 37 °C for 24 h, resulting in a visible gel at the bottom of the PCR tube. After transcription, (1) 20 µL of IVT-buffer (control) or (2) 20 µL of separately transcribed HHRz (overnight, 37 °C) were added. Samples were mixed by pipetting, ncubated for 4 h at 37 °C with shaking at 700 rpm, and then left overnight at 37 °C without shaking. b) PCR tubes after 22 h of incubation. The control sample (1) retains the visible RNA gel, while the HHRz-treated sample (2) In the control (1), the gel remains intact and can be extracted as a single piece with a pipette, indicating that degradation only occurs in the presence of the trans-acting HHRz.

#### 2.5 Figure S5: Denaturing polyacrylamide gel electrophoresis of hairpin ribozyme cleavage products

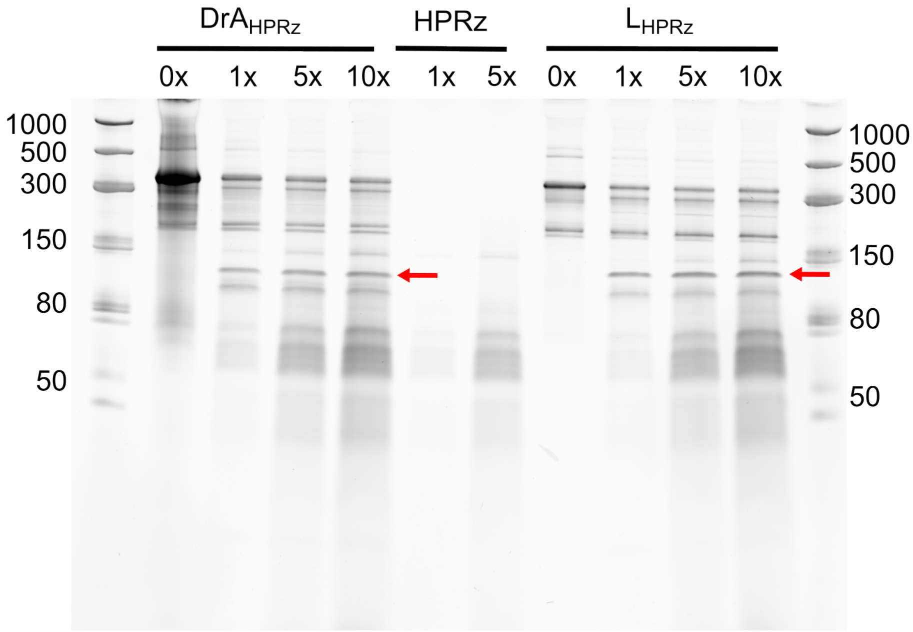

Figure S5: Denaturing polyacrylamide gel electrophoresis of hairpin ribozyme cleavage products. *In vitro* transcribed RNA nanostars (corresponding design is written on top of the gel) were incubated with increasing volumes of ribozyme (1×, 5×, 10×), where 1× corresponds to 1 µL of ribozyme added to 1 µL of target RNA in a total volume of 50 µL. Lanes labeled 0× serve as cleavage-negative controls with no ribozyme added, the lanes labeled HHRz or HPRz 1× and 5× represent ribozyme-only controls, where ribozymes were diluted (1 µL or 5 µL )in 50 µL IVT-buffer without droplet RNA (for more details see methods). Cleavage products were resolved by 10 % denaturing PAGE stained with GelRed®. Low Range ssRNA Ladder is used as reference, length in bases is reported as text close to the corresponding bands. Red arrows indicate expected cleavage products. Cleavage of full-length constructs $DrA_{HPRz}$/ $L_{HPRz}$ (272/274 nt) is expected to yield three fragments with a lenght of 23 nt, 135/137 nt and 114 nt.

#### 2.6 Figure S6: Additional replicate for DrA$_{\mathbf{HPRz}}$

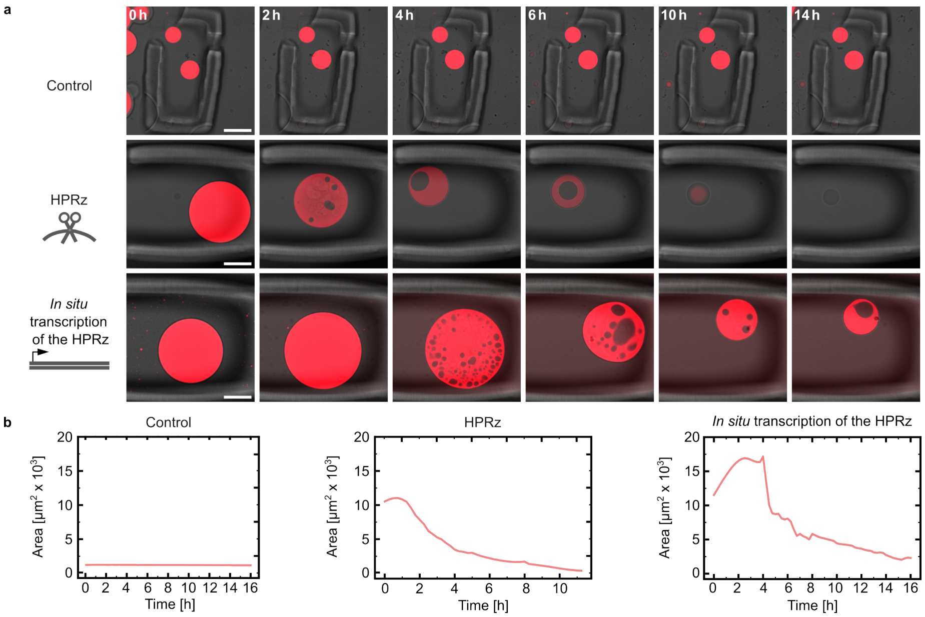

Figure S6: Additional replicates for DrA$_{\mathrm{HPRz}}$. Confocal time series using the HPRz for dissolution of DrA$_{\mathrm{HPRz}}$. **a)** Confocal timeseries (fluorescence and brightfield overlay, droplets contain MGA and are labeled with malachite green dye, $\lambda_{ex} = 640\,\mathrm{nm}$) for a technical replicate of dissolving RNA droplets using the HPRz. At $t = 0$, buffer (top), overnight transcribed ribozyme (middle) or IVT containing the DNA template for the HPRz, transcribing the ribozyme *in situ* (bottom) was added. Scale bars:50 µm. **b)** Droplet area over time for the droplets shown in a)

#### 2.7 Figure S7: Quantification of vacuole-to-area ratio in $DrA_{HPRz}$

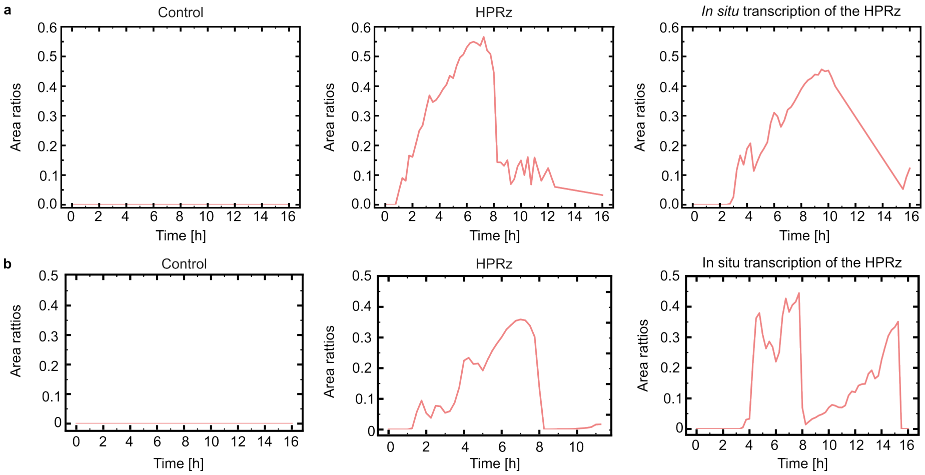

Figure S7: Quantification of vacuole-to-area ratio in $DrA_{HPRz}$. Transient vacuole formation observed and quantified for **a)** the droplets in Figure 3b and **b)** for the droplets in Figure S6. These vacuoles form and disappear over the observation time of 16 h. The droplet detection and vacuole-to-droplet area measurements were performed using a custom-written python script.

#### 2.8 Figure S8: Effect of cleavage on droplet size for $DrA_{HPRz}$

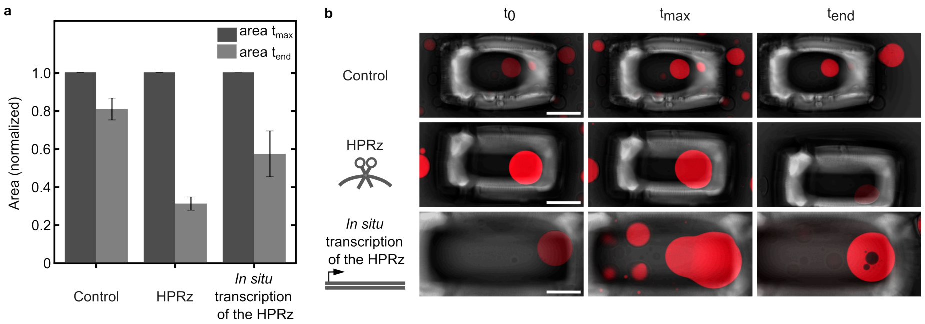

Figure S8: Effect of cleavage on droplet size for $DrA_{HPRz}$. HPRz was used for dissolution of $DrA_{HPRz}$. **a)** Comparing the normalized area of the droplets at the timepoint when they reach their maximal diameter to the droplet area at the end of the timelapse (n= 6 regions). Data for the plot was pooled from one replicate. The droplet detection and area measurements were performed using a custom-written python script. **b)** Confocal micrographs (fluorescence and brightfield overlay, droplets contain MGA and are labeled with malachite green dye, $\lambda_{ex} = 640\,\text{nm}$) for one biological replicate of dissolving RNA droplets using the HPRz. One of the regions (per condition) plotted in a) is shown at the start of the timelapse ($t_0$), at the time where they reach their maximal size ($t_{max}$) and at the end of the timelapse ($t_{end}$), ending 22 h and 15 min after the addition of the HPRz. Scale bars: 50 µm.

#### 2.9 Figure S9: Additional replicates characterizing the diffusion of fluorescently labeled hairpin ribozyme into the droplet

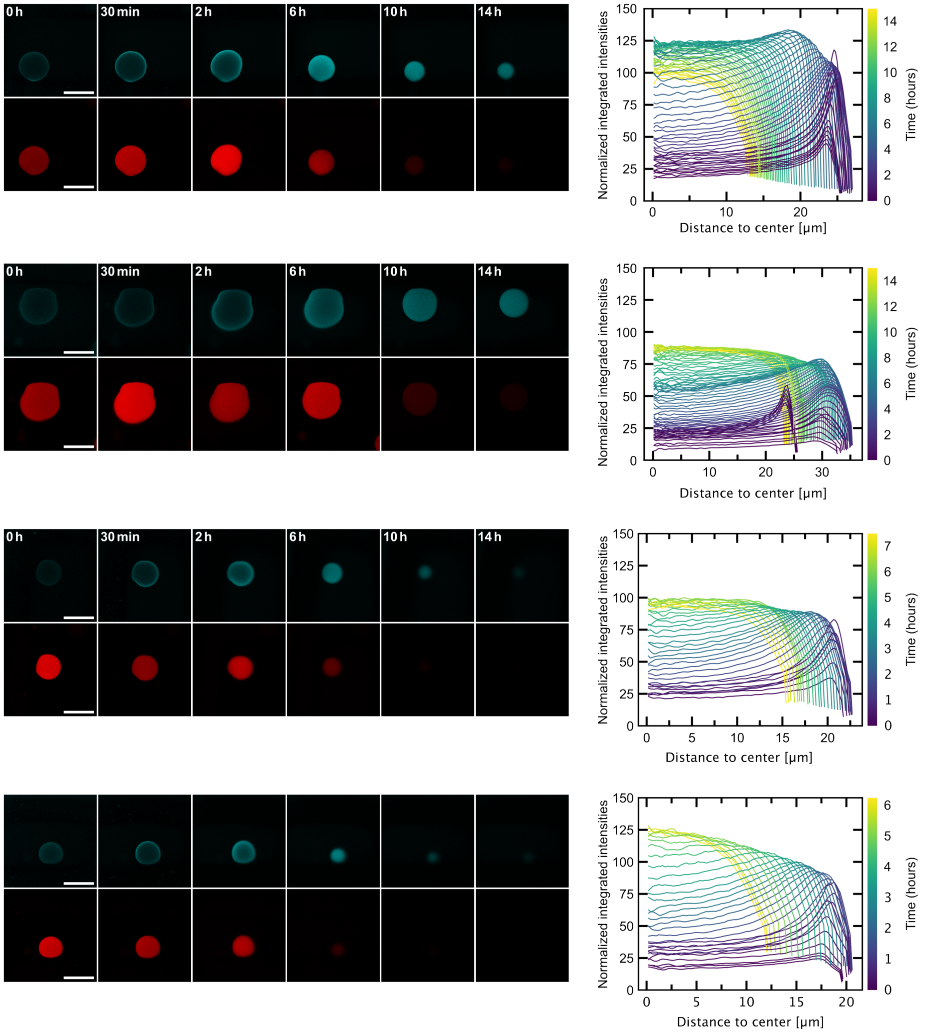

Figure S9: Confocal timeseries of caged RNA droplets (left) after the addition of fluorescently labeled HPRz. At $t = 0$, the HPRz was flushed in. The HPRz (blue) is labeled with fluorescein, $\lambda_{ex} = 488$ nm. Radial intensity profiles (right) showing the diffusion of fluorescently labeled HPRz into a single RNA droplet over time. Normalized integrated intensities were measured as a function of distance from the droplet center. The color gradient represents different time points over a 14 h period or until the droplet is dissolved. Scale bars: 50 µm.

#### 2.10 Figure S10: Ribozyme cleavage is sequence-specific

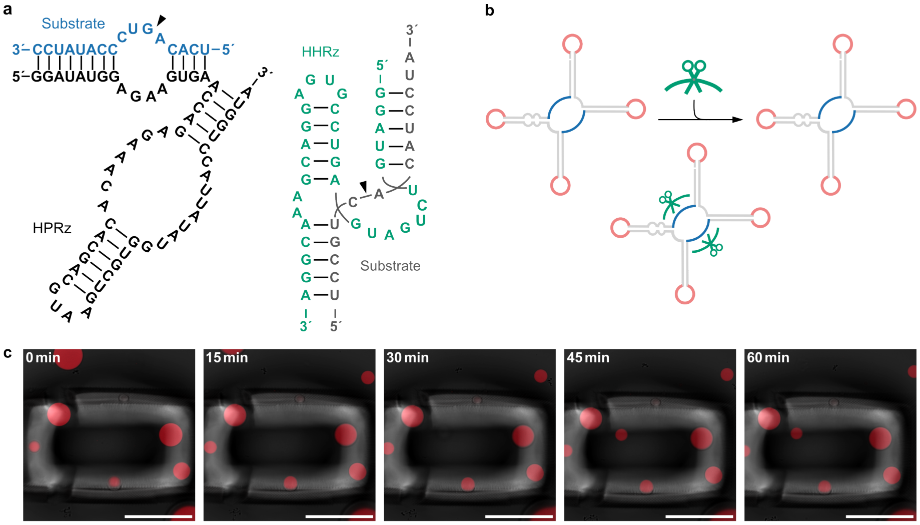

Figure S10: Ribozyme cleavage is sequence-specific. Verification of the sequence specificity of ribozyme cleavage by adding the HHrz to $\mathrm{DrA_{HPRz}}$ **a)** RNA sequence of the HPRz (pink) and the HHRz (green) used in this experiment. **b)** Schematic representation illustrating the incorporation of the substrate sequence from the HPRz into the nanostar design. Addition of the HHRz ribozyme should not lead to cleavage, since the wrong substrate sequence is present. **c)** Confocal timeseries (fluorescence and brightfield overlay, droplets contain MGA and are labeled with malachite green dye, $\lambda_{ex} = 640\,\mathrm{nm}$) of $\mathrm{DrA_{HPRz}}$. At $t = 0$, the HHRz was added to the droplets. Scale bars: 50 µm.

#### 2.11 Figure S11: Titrating linker ratios

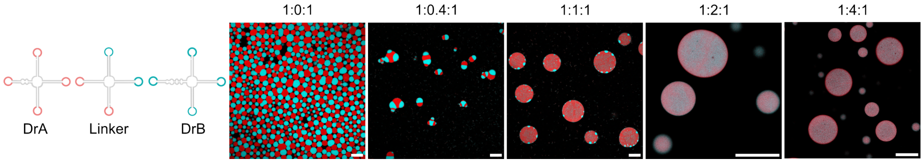

Figure S11: Titrating linker ratios. Confocal micrographs of mixed RNA droplets using different DNA template ratios for DrA: L: DrB. Red droplets contain MGA and are labeled with malachite green dye, $\lambda_{ex} = 640\,\text{nm}$, blue droplets contain BrA and are labeled with DFHBI-1T, $\lambda_{ex} = 488\,\text{nm}$. Images were acquired more than 24 h after the start of transcription. Scale bars: 50 µm.

#### 2.12 Figure S12: Additional replicates of ribozyme-induced segregation of RNA droplets

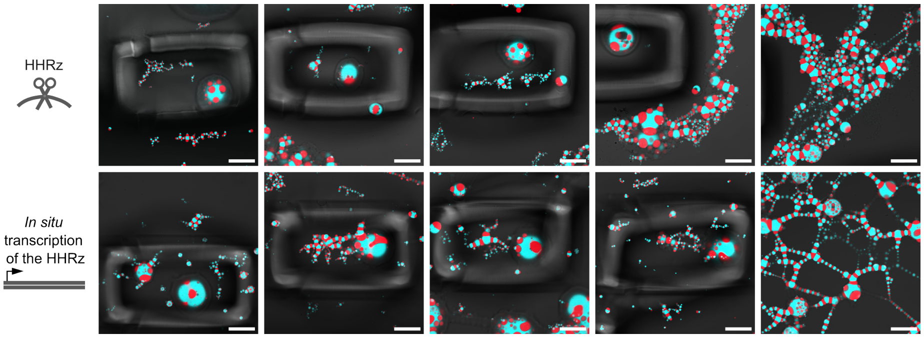

Figure S12: Confocal micrographs (fluorescence and brightfield overlay, red droplets contain MGA and are labeled with malachite green dye, $\lambda_{ex} = 640\,\text{nm}$, blue droplets contain BrA and are labeled with DFHBI-1T, $\lambda_{ex} = 488\,\text{nm}$) acquired 15 h after addition of the HHRz, as shown in Figure 4b. The focal plane was changed to show smaller droplets that were not in focus in the caged region as well as selecting other regions to show uncaged droplets (right). Scale bars: 50 µm.

#### 2.13 Figure S13: Comparing the ribozyme cleavage efficiency to enzymatic cleavage using RNase H

To compare ribozyme cleavage with enzymatic strategies, we selected the endonuclease RNase H, which specifically degrades RNA when hybridized to a complementary DNA strand in an RNA/DNA heteroduplex [1]. DrA, DrB and a linker containing two single-stranded regions (16 nt, $L_{HPRz}$ were transcribed using a 1:1:2 DNA template ratio. Upon addition of two complementary DNA strands ($DNA_{c1}$ and $DNA_{c2}$), the single-stranded regions in the linker form an RNA-DNA duplex, which are subsequently cleaved by RNase H (Figure S13a).

The segregation behavior closely resembled that observed with ribozyme cleavage: over the course of several hours the droplets first collapse, accompanied by hole formation in the center of the droplet. Subsequently, the monomers of DrA and DrB rearrange followed by their segregation. To confirm that RNase H cleavage was sequence-specific, we performed two control experiments: one without the addition of complementary DNA strands and another with complementary DNA but without RNase H. In both cases, segregation did not occur (Figure S13b).

To quantitatively describe the segregation kinetics, we also determined the Pearson's correlation coefficient ($R_r$) over time. Similar to the ribozyme-mediated cleavage $R_r$ stabilizes around $-0.5$ , but with strong fluctuations. However, the initial phase of segregation was not captured, as the necessary steps: gluing the channels, transferring the sample to the confocal microscope, and setting up the tile regions for the timelapse took approximately 20 min. To obtain a precise $t_0$ corresponding to the time of addition, we would need to use the microfluidic pump in this setup as well.

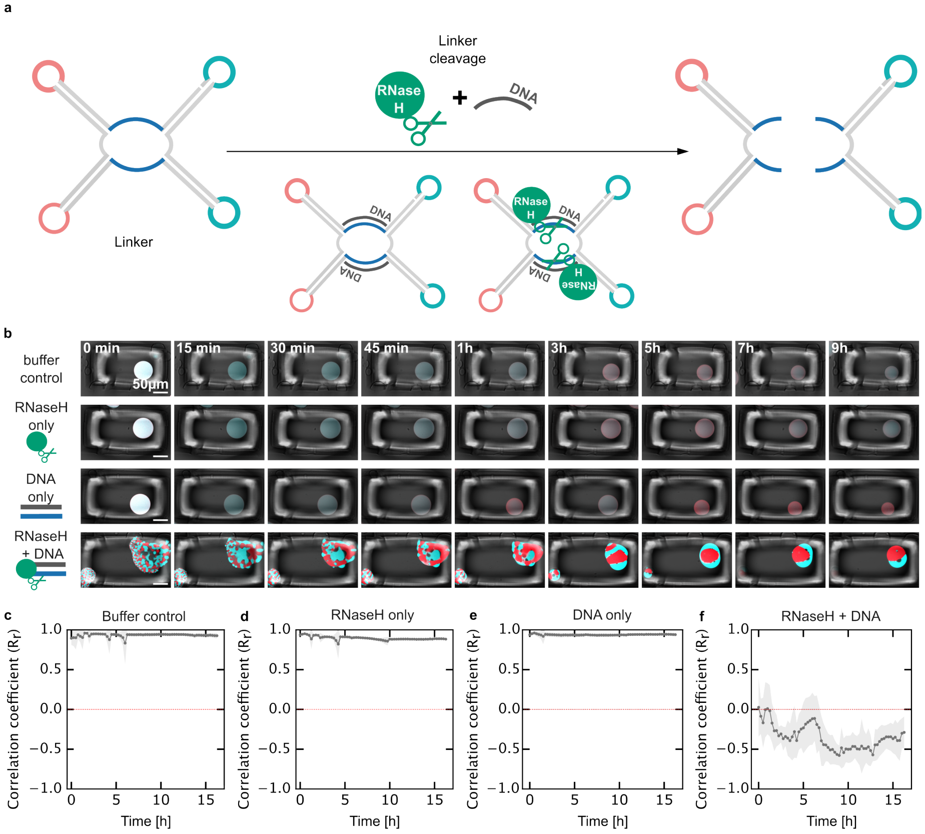

Figure S13: Comparing the ribozyme cleavage efficiency to enzymatic cleavage using RNase H. **a)** Schematic representation exemplifying linker cleavage using RNase H. Upon addition of two complementary DNA strands, the single-stranded regions in the linker form an RNA-DNA duplex, which is subsequently cleaved by RNase H. **b)** Confocal timeseries of caged RNA droplets (fluorescence and brightfield overlay, red roplets contain MGA and are labeled with malachite green dye, $\lambda_{ex} = 640\,\text{nm}$, blue droplets contain BrA and are labeled with DFHBI-1T, $\lambda_{ex} = 488\,\text{nm}$). $t = 0$ corresponds to the start of the timelapse about 20 min after the addition of either 1x RNase H buffer, RNase H only, DNA only or RNase H and DNA. Scale bars: 50 µm. **c)**, **d)**,**e)**, **f)** Colocalization analysis (Pearson correlation) was performed and Pearson $R_r$ values are plotted over time (mean ± s.d., n = 3 regions for **c)**, **d)**, **f)**, n = 2 regions for **e)**).

#### 2.14 Figure S14: Mixed droplets are stable for days

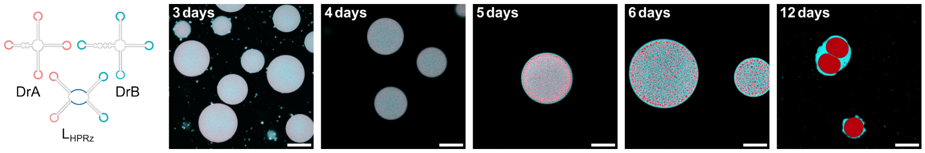

Figure S14: Mixed droplets are stable for days. Confocal micrographs of mixed RNA droplets (DrA, DrB and the respective linker L) monitoring their stability over several days. Red droplets contain MGA and are labeled with malachite green dye, $\lambda_{ex} = 640\,\text{nm}$, blue droplets contain BrA and are labeled with DFHBI-1T, $\lambda_{ex} = 488\,\text{nm}$. DrA, DrB and $L_{\text{HPRz}}$ were transcribed together with a DNA template ratio of 1:1:2 and incubated at 37 °C. The onset of segregation without ribozyme-catalyzed linker cleavage can only be observed after 5 days and is clearly evident after 12 days. This is likely due to non-specific degradation of the single stranded regions in the linker and the BrA in DrB. Scale bars: 50 µm

#### 2.15 Figure S15: Agarose gel electrophoresis of PCR amplified DNA templates.

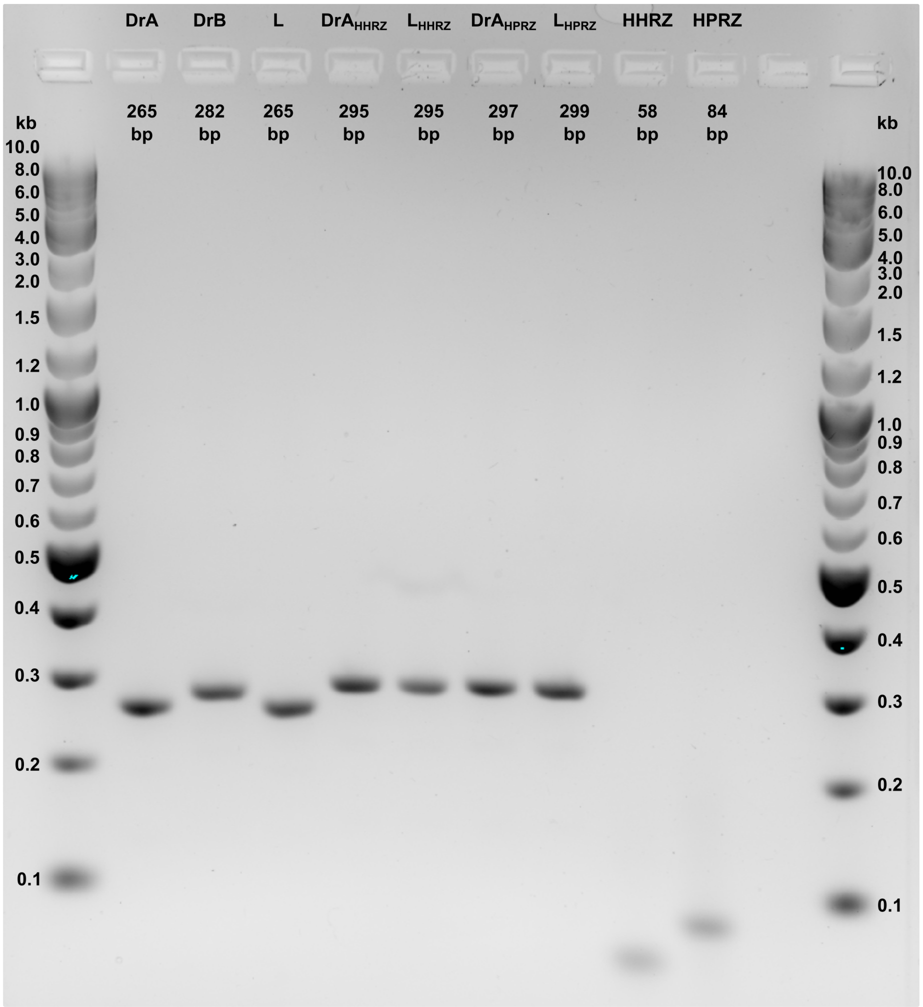

Figure S15: Agarose gel electrophoresis of PCR products. 3% argarose gel was run at 90 V and stained with Midori Green Advance DNA Stain. TriDye® 1 kb Plus DNA Ladder is used as reference, length in kilo base pairs is reported as text close to the corresponding bands. Visual inspection of the gel confirms the expected sizes of the DNA templates (expected length in base pairs (bp) is reported on top

### 3 Supplementary Videos

#### 3.1 Video S1

Confocal timelapse of caged RNA droplets ($DrA_{HPRz}$), showing that the droplet remains inside of the cage for 22 h. The cage prevents movement and fusion with other droplets and therefore allows to observe individual droplets over time. Droplets contain MGA and are labeled with malachite green dye, $\lambda_{ex} = 640$ nm, fluorescence and brightfield overlay. From the starting point up to 22 h and 15 min, every frame is separated by a 15 min time interval. Timestamps are shown on the top left.

#### 3.2 Video S2

Confocal timelapse of caged RNA droplet ($DrA_{HHRz}$), dissolving upon addition of the HHRz. Scale bar: 50 µm. Droplet contains MGA and is labeled with malachite green dye, $\lambda_{ex} = 640$ nm, fluorescence and brightfield overlay. From the starting point up to 47 min, every frame is separated by a 30 s time interval. Timestamps are shown on the top left.

#### 3.3 Video S3

Confocal timelapse of caged RNA droplet ($DrA_{HPRz}$), dissolving upon addition of HPRz. Scale bar: 50 µm. Droplet contains MGA and is labeled with malachite green dye, $\lambda_{ex} = 640$ nm, fluorescence and brightfield overlay. From the starting point up to 16 h, every frame is separated by a 15 min time interval. Timestamps are shown on the top left.

#### 3.4 Video S4

Confocal timelapse of caged RNA droplet ($DrA_{HPRz}$), dissolving upon addition of the IVT mixture containing the DNA template for the HPRz, transcribing the ribozyme *in situ*. Scale bars: 50 µm. Droplet contains MGA and is labeled with malachite green dye, $\lambda_{ex} = 640$ nm, fluorescence and brightfield overlay. From the starting point up to 16 h, every frame is separated by a 15 min time interval. Timestamps are shown on the top left.

#### 3.5 Video S5

Confocal timelapse of mixed, caged RNA droplets (DrA, DrB, $L_{HHRz}$) segregating upon addition of the transcribed HHRz. Red droplets contain malachite green aptamer (MGA) and are labeled with malachite green dye, $\lambda_{ex} = 640$ nm, blue droplets contain BrA and are labeled with DFHBI-1T, $\lambda_{ex} = 488$ nm, droplets are mixed in the beginning using a ratio of 1:2:1 (DrA:L:DrB). From the starting point up to 14 min 15 s, every frame is separated by a 45 s time interval followed by a short 80 s pause to seal the chamber. Imaging then continued until 1 h 4 min 5 s, with a 45 s time interval, after which the interval was extended to 10 min or overnight acquisition. Timestamps are shown on the top left.

### References

1. Cerritelli, S. M. & Crouch, R. J. Ribonuclease H: the enzymes in eukaryotes. *The FEBS journal* **276,** 1494–1505 (2009).

### 4 Uncropped and non-inverted gel images

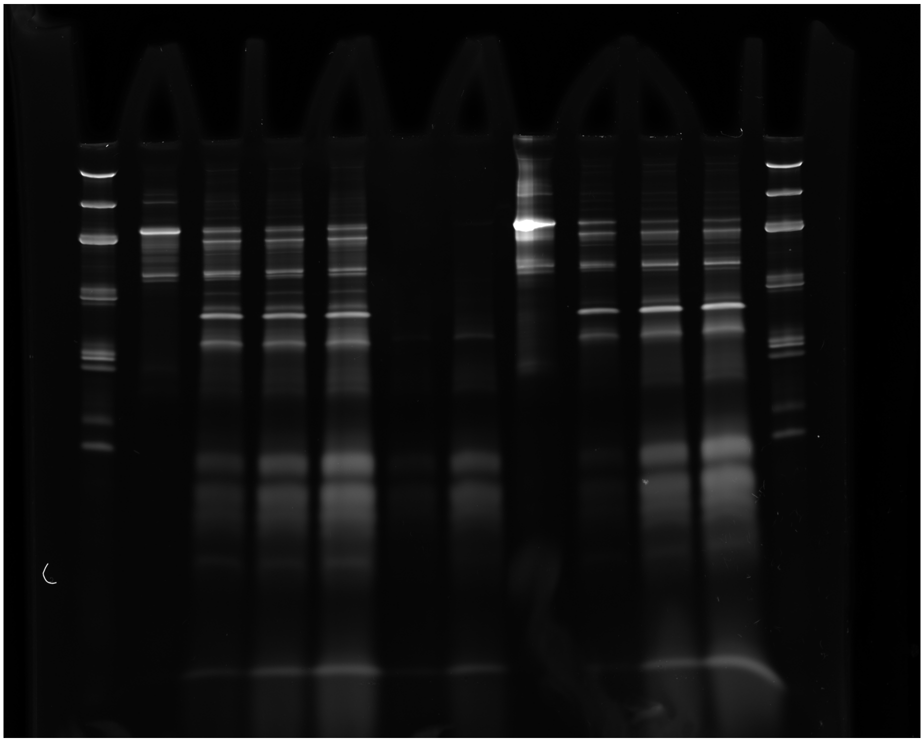

Figure S16: Uncropped and non-inverted PAGE of cleavage products. Cleavage using the HHRz, Figure S4.

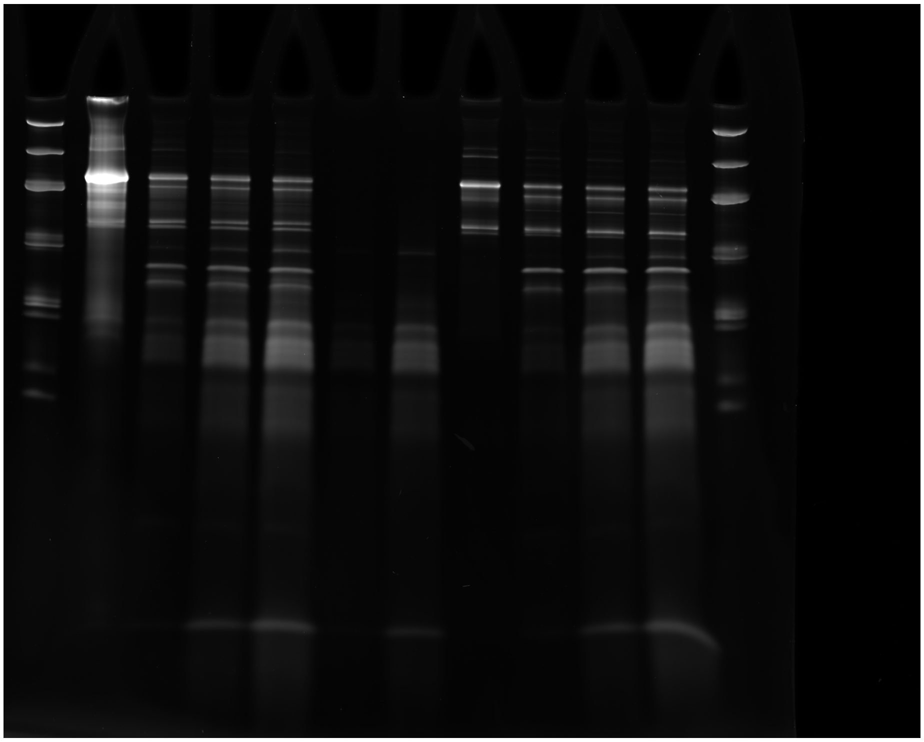

Figure S17: Uncropped and non-inverted PAGE of cleavage products. Cleavage using the HPRz, Figure S5.

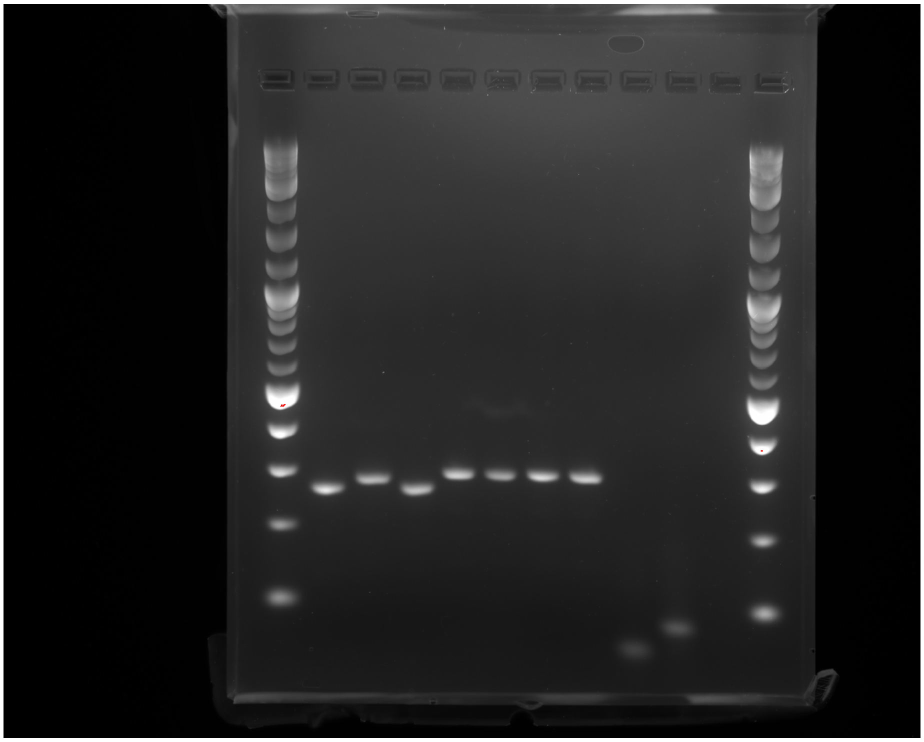

Figure S18: Uncropped and non-inverted agarose gel electrophoresis of PCR products, Figure S15
